# Creation and analysis of enzymatically active micrometer protein-capsules

**DOI:** 10.1101/2020.10.22.350488

**Authors:** Kai Melvin Schakowski, Christian Elm, Jürgen Linders, Michael Kirsch

**Affiliations:** Institute of Physiological Chemistry, University of Duisburg-Essen, University Hospital Essen, Hufelandstraße 55, 45147 Essen, Germany; Department of Physical Chemistry, University of Duisburg-Essen, Universitätsstraße 5, 45141 Essen, Germany; Center for Nanointegration Duisburg-Essen (CeNIDE), University of Duisburg-Essen, Carl-Benz-Straße 199, 47057 Duisburg, Germany

**Keywords:** catalase, albumin, oxygen, microencapsulation, microcapsules, enzymes

## Abstract

This work describes a general method for the encapsulation of enzymes with albumin as wall material and the enzyme catalase as prime example. Care was taken for the preparation of biochemically active sub-micrometer particles in order to prevent oxygen toxicity induced by artificial oxygen carriers of any type. In cell culture experiments, capsules containing catalase did not exhibit any harmful activities in the absence of peroxides. In the presence of hydrogen peroxide application of low and medium dosed capsules below 0.05 vol% (final concentration 0.001 vol%) even increased the cell damaging process. However, a higher dosage of capsules (> 0.05 vol%) prevented completely cellular disruption induced by 5 mM hydrogen peroxide and decreased up to 90% of cellular damage at higher peroxide concentrations. These results demonstrated that encapsulated catalase was enzymatically active and the over-all activity of prepared catalase capsules was determined to be > 1000 U ∙ mL^−1^ ∙ vol%^−1^.

## 1 Introduction

In order to counteract the rising shortness on blood donations all over the world, the development of artificial blood substitutes has been investigated with several different approaches (Müller *et al.* 2015, Chung *et al.* 2016, Ellingson *et al.* 2017). Possible blood substitutes were created using either hemoglobine-based oxygen carriers (HbOCs) (Xiong *et al.* 2012, Schakowski *et al.* 2020) or perfluorocarbon-based oxygen carriers (PFCOCs) (Wrobeln *et al.* 2017). HbOCs as well as PFCOCs come with a shell of either biodegradable polymers or albumin (Chang and Yu 1995, Bauer *et al.* 2010, Xiong *et al.* 2012) whereas free hemoglobin has widely been ruled out due to nephrotoxicity (Simoni *et al.* 1997).

Promisingly, all these approaches have the drawback, that more intracellular components, e.g. enzymes or co-factors, are needed for the adequate replacement of red blood cells (RBCs). In order to reduce damage induced by free radicals, RBCs naturally contain various additional enzymes, such as catalase and superoxide dismutase. In 1984 TURRENS has shown that encapsulation of RBC-enzymes into liposomes is principally possible (Turrens *et al.* 1984). In this work we prove that the encapsulation of RBC-enzymes is possible with the more modern method of encapsulation into a shell of albumin. We take advantage of organic, inorganic and physical chemistry for the preparation of biochemically active sub-micrometer particles to prevent oxygen toxicity induced by artificial oxygen carriers of any type.

In 1913 Heard already proposed a coherence between metal cations, carbonates and the precipitation of proteins (Heard 1913). In this work, the co-precipitation of catalase together with the hardly soluble MnCO_3_ template was used according to a protocol introduced by Johnson (Johnson 1982). The subsequent addition of BSA to these MnCO_3_-catalase generated an albumin wall thus leading to particles consisting of a MnCO_3_-catalase core and a BSA shell.

## 2 Experimental

### 2.1 Materials

Bovine serum albumin fraction V (BSA), Manganese(II) chloride tetrahydrate, Ethylenediaminetetraacetic acid disodium salt dihydrate (Na_2_EDTA) and Triton X-100 were purchased from Sigma-Aldrich (Darmstadt, Germany); Sodium carbonate anhydrous and Sodium hydroxide were purchased from Merck (Darmstadt, Germany); Disodium phosphate dehydrate was purchased from Carl Roth (Karlsruhe, Germany); Ringer’s solution was purchased from Fresenius Kabi (Bad Homburg, Germany); Gibco Minimum Essential Medium (MEM), Gibco Fetal bovine serum (FBS), Gibco Pen Strep and Gibco 0.05% Trypsin-EDTA were purchased from Thermo Fisher Scientific (Schwerte, Germany); Catalase from bovine liver was purchased from SERVA (Heidelberg, Germany); Genipin was purchased from TCI Chemicals (Eschborn, Germany). All materials were of highest quality available.

### 2.2 Synthesis of Albumin-Catalase capsules (ACCs)

ACCs were created by a method first presented by XIONG *et al.* (Xiong *et al.* 2012, 2013, Li *et al.* 2017) and later modified by SCHAKOWSKI *et al.* (Schakowski *et al.* 2020). Cross-linking of the BSA molecules by genipin as described in previous works (Butler *et al.* 2003, Yoo *et al.* 2011, Shahgholian *et al.* 2017, Schakowski *et al.* 2020) results in a stable shell of BSA around the core particle. Dissolution of MnCO_3_ from the core by EDTA finally leads to the release of formerly entrapped catalase to the inside of the created capsule.

Two aqueous solutions of 25 mL, each containing 10 mg ∙ mL^−1^ Catalase, 1 mg ∙ mL^−1^ BSA and either 0.25 mM Na_2_CO_3_ or 0.25 mM MnCl_2_, respectively, were prepared. The Na_2_CO_3_ solution was placed in a 100 mL Erlenmeyer flask and the MnCl_2_ solution was added slowly under continuous vigorous stirring. After total merging of the two solutions, stirring was continued for two more minutes. Subsequently 125 mg BSA was carefully added in portions and after total dissolution of the BSA, stirring was continued for five more minutes. The resulting suspension was equally split into two 50 mL reaction vessels and centrifuged at 1,000 g for two minutes. The supernatant was discarded and the residue was re-suspended in 40 mL of Ringer’s solution each using shaking and ultrasonic bath. Centrifugation and washing were repeated two more times, before the residue was re-suspended in 40 mL of Milli-Q water each. The resulting suspensions were then merged into a 200 mL Erlenmeyer flask and under continuous stirring 20 mL of 940 μM Genipin solution in Milli-Q water were added dropwise, resulting in a final concentration of 188 μM Genipin. The aperture of the Erlenmeyer flask was then covered by aluminium foil and the suspension was stirred for 24 hours at room temperature.

The suspension was equally split into two new 50 mL reaction vessels and again centrifuged for two minutes at 1,000 g before the supernatants were discarded. 40 mL of Na_2_EDTA solution of pH 7.4 were added to each reaction vessel and the closed reaction vessels were shaken and kept in ultrasonic bath until all residue had fully dissolved. After centrifugation for two minutes at 1,500 g and discarding of the supernatants, this step was repeated once more. Emerging foam on the top of the supernatants was skimmed off and residues were suspended in 1 mL of Ringer’s solution before being merged into a new suitable reaction vessel.

Prepared ACCs can be stored at 4 °C or −20°C in Ringer’s solution for more than six months maintaining a useful level of enzymatic activity.

This procedure has also been performed using ZnCl_2_ instead of MnCl_2_.

### 2.3 Characterization

#### 2.3.1 Transmission Electron Microscopy (TEM)

Transmission Electron Microscopy was performed using a JEM 1400 Plus TEM (JEOL, Freising, Germany).

#### 2.3.2 Respirometry assay

Respirometry was performed on an Oroboros Oxygraph-2k Fluorespirometer (Oroboros Instruments, Innsbruck, Austria).

The chamber volume *V* of the fluorespirometer was adjusted to 2 mL and 37 °C and filled with 50 mM sodium phosphate buffer or artificial serum, both of pH 7.4. After the oxygen background of the fluorespirometer had stabilized, 25 μL of freshly prepared 120 mM H_2_O_2_ in 50 mM sodium phosphate buffer of pH 7.4 were injected into the chamber using a Hamilton syringe. After the oxygen flux had stabilized due to accelerated auto-degradation of H_2_O_2_ in artificial serum, 50 μL of ACC-solution of different concentrations *c* was injected into the chamber and oxygen flux was measured for five minutes.

The oxygen flux was corrected for the background flux due to auto-degradation and the average corrected oxygen flux *f* was obtained by linear regression. The enzymatic activity *A* was calculated as A = (2f ∙ V) / (c ∙ 50 μL), taking into account the ratio of produced O_2_ and degraded H_2_O_2_ in 2 H_2_O_2_ → O_2_ + 2 H_2_O.

The artificial serum (AS) was created by dissolving 298.2 mg (4 mmol) KCl, 162.6 mg (0.8 mmol) MgCl_2_ ∙ 6 H_2_O, 266.4 mg (2.4 mmol) CaCl_2_, 2016.2 mg (24 mmol) NaHCO_3_, 89.7 mg (0.65 mmol) NaH_2_PO_4_ ∙ H_2_O, 92.3 mg (0.65 mmol) Na_2_HPO_4_, 6665.1 mg (114.05 mmol) NaCl and 50 g BSA in 1000 mL MilliQ water. The pH was adjusted to 7.4 at 37 °C using NaOH and HCl, each ≤ 1 M. AS was split into portions of 50 mL and stored at −20 °C. AS once thawed should be kept at 37 °C and should be not be refrozen.

#### 2.3.3 Cell culture

Cells were cultured in minimum essential medium (MEM) Eagle supplemented with 25 mM sodium bicarbonate, 10% FBS, 2 mM L-glutamine, and 100 units/mL penicillin G and 0.1 mg/mL streptomycin in 75 cm^2^ plastic flasks at 37 °C in a humidified atmosphere of 95% air and 5% CO_2_. Subcultures were obtained by trypsinisation (0.05% trypsin-EDTA). For experimental use, cells were transferred to 6-well plates.

##### 2.3.3.1 Cytotoxicity assays

The standard cytotoxicity model of murine fibroblast-derived cell line L929 (American Type Culture Collection, NCTC clone 929 of strain L) was used for the experiments (Lomonosova *et al.* 1998). Cells were incubated in 6-well-plates under the described conditions over night. The medium was removed and 2 mL of 50 mM sodium-phosphate buffer of pH 7.4 and 50 μL of trypan blue solution were added.

For cytotoxicity of H_2_O_2_, 50 μL of freshly prepared H_2_O_2_-solution was added, so that final H_2_O_2_-concentrations in the wells were 0, 1.25, 2.5, 5.0, 7.5 and 10 mM. Cells were counted by hand after 0, 30, 60, 90, 120, 150 and 180 minutes using an inverted microscope and the fraction of cells that had taken up trypan blue as a marker of loss of membrane integrity was calculated. For the duration of the experiment, cells were kept under described conditions.

For the cytotoxicity assay of ACCs, 100 μL of ACC-suspension of different concentrations was added instead of H_2_O_2_. The period of observation was prolonged to 24 hours.

##### 2.3.3.2 Protection assay

To determine the protective capacity of prepared ACCs, cells were incubated in 6-well-plates under described conditions. Again 50 μL of trypan blue solution were added. Then 50 μL of ACC-suspension were added with original concentrations of 0.001, 0.005, 0.010, 0.100, 0.250 and 0.500 vol%.

Results from H_2_O_2_ cytotoxicity assays served as control.

## 3 Results and Discussion

### Physical properties

Transmission electron microscopy was used to determine shape and size of the ACCs. Measuring the longest diameter of 452 single ACCs, prepared by using a MnCO_3_ template, yielded in a mean diameter of 1.245 μm with a standard deviation of 0.272 μm. The overall range of diameters ranges from 0.433 μm to 1.951 μm. As seen in figure 2, particles are shaped ellipsoid rather than round, whereas the core displays to have a peanut-like shape.

**Figure 1:**
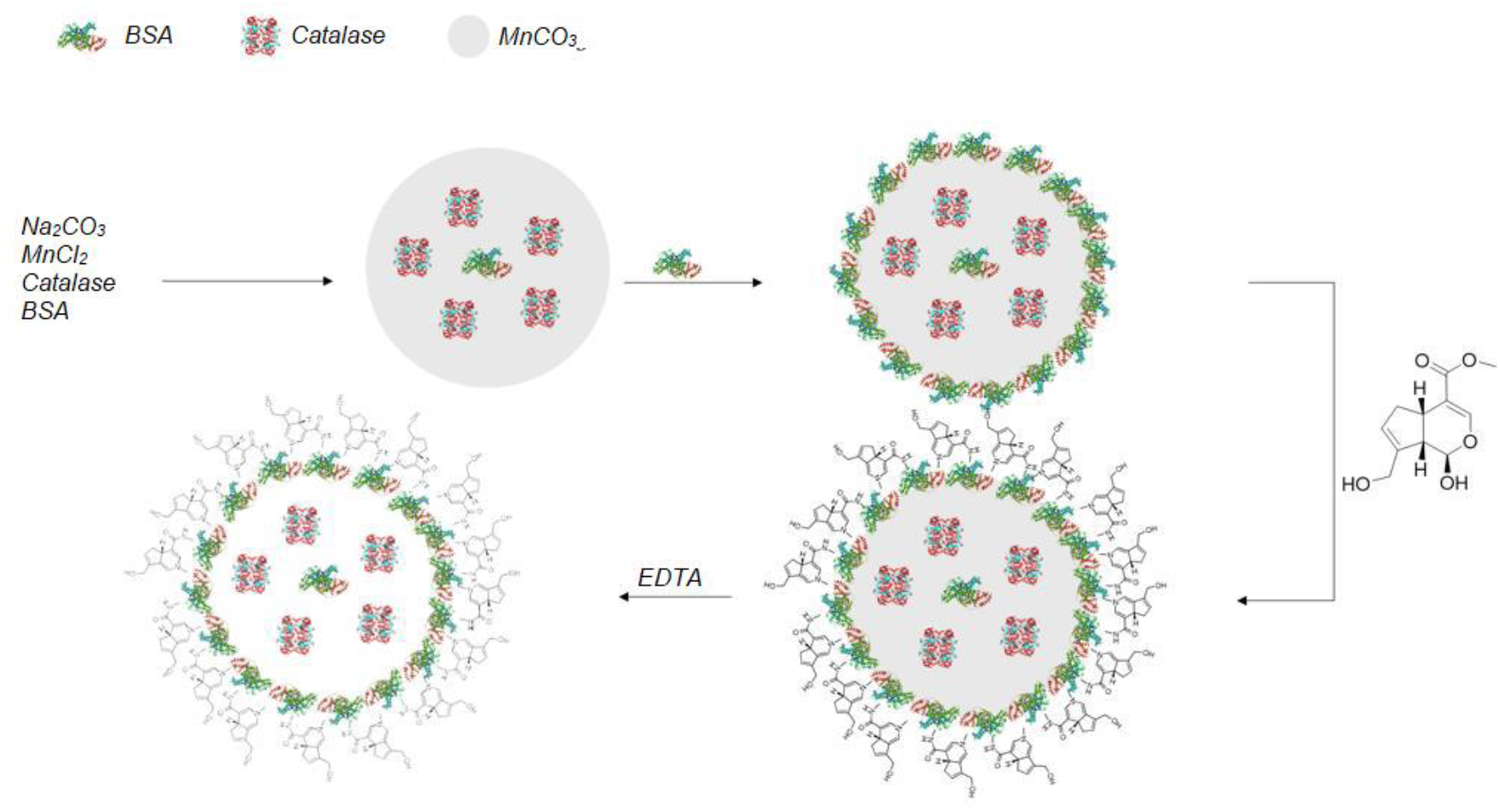
Synthesis of ACCs by co-precipitation of MnCO_3_ and catalase, followed by adsorption of BSA and cross-linking by genipin. The MnCO_3_ core is then dissolved using EDTA.

**Figure 2:**
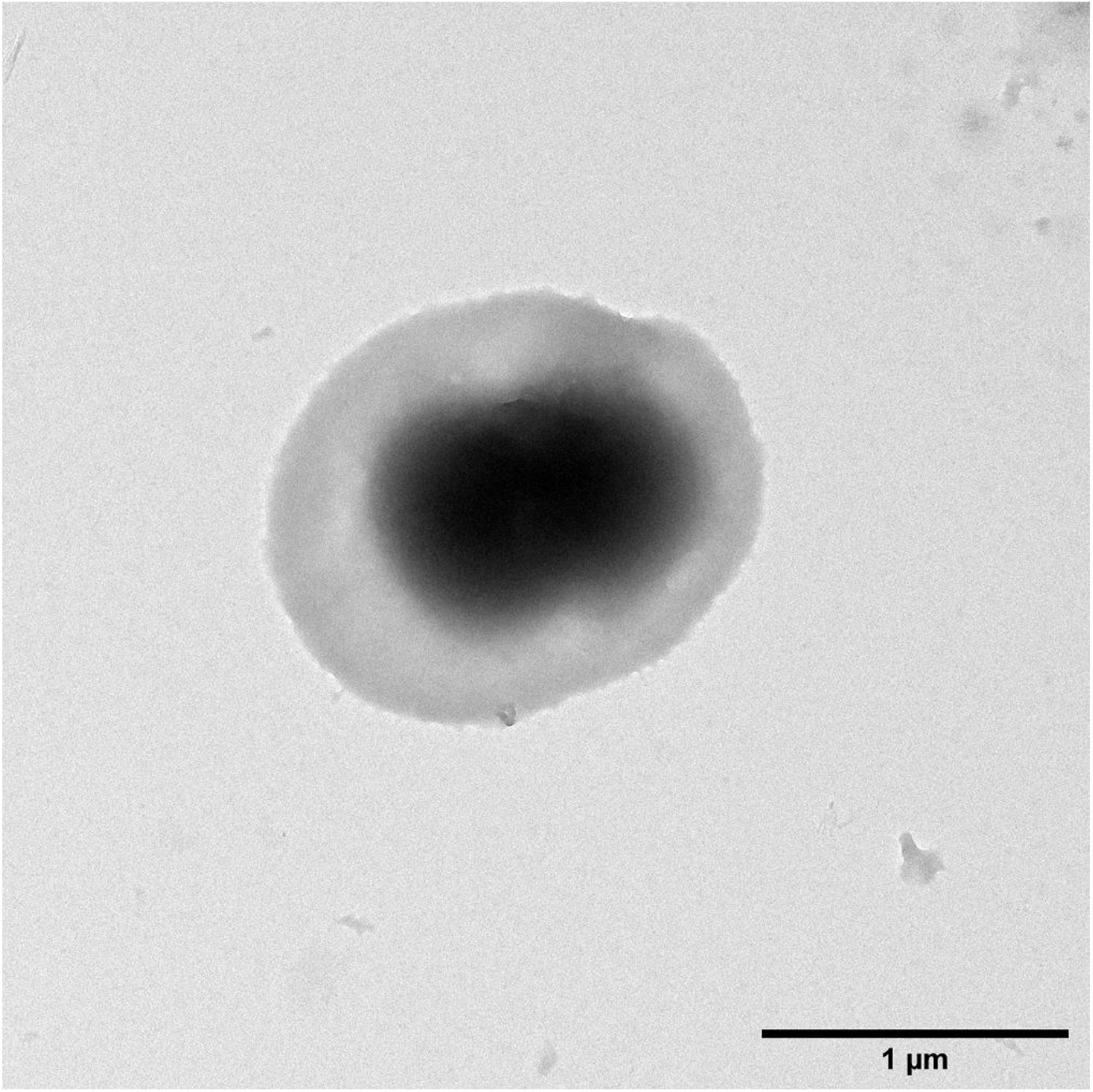
A solitary ACC from MnCO3 template pictured by TEM. Its longest diameter measures about 1.65 μm.

ζ-potential was determined to be −30.64 mV ± 7.76 mV. Each sample was measured three times and the overall mean and standard deviation were calculated.

Using ZnCO_3_ as a template, solitary capsules observed were about 20% smaller, as seen in figure 3 (left), but have the great disadvantage of forming huge bulky adducts with large amount of inter-capsular cross-linking (right). Unfortunately, the changing of various parameters (e.g. kind of enzyme, albumin and cross-linking agent) failed in decreasing the amount of inter-capsular cross-linking to insignificant yields.

**Figure 3:**
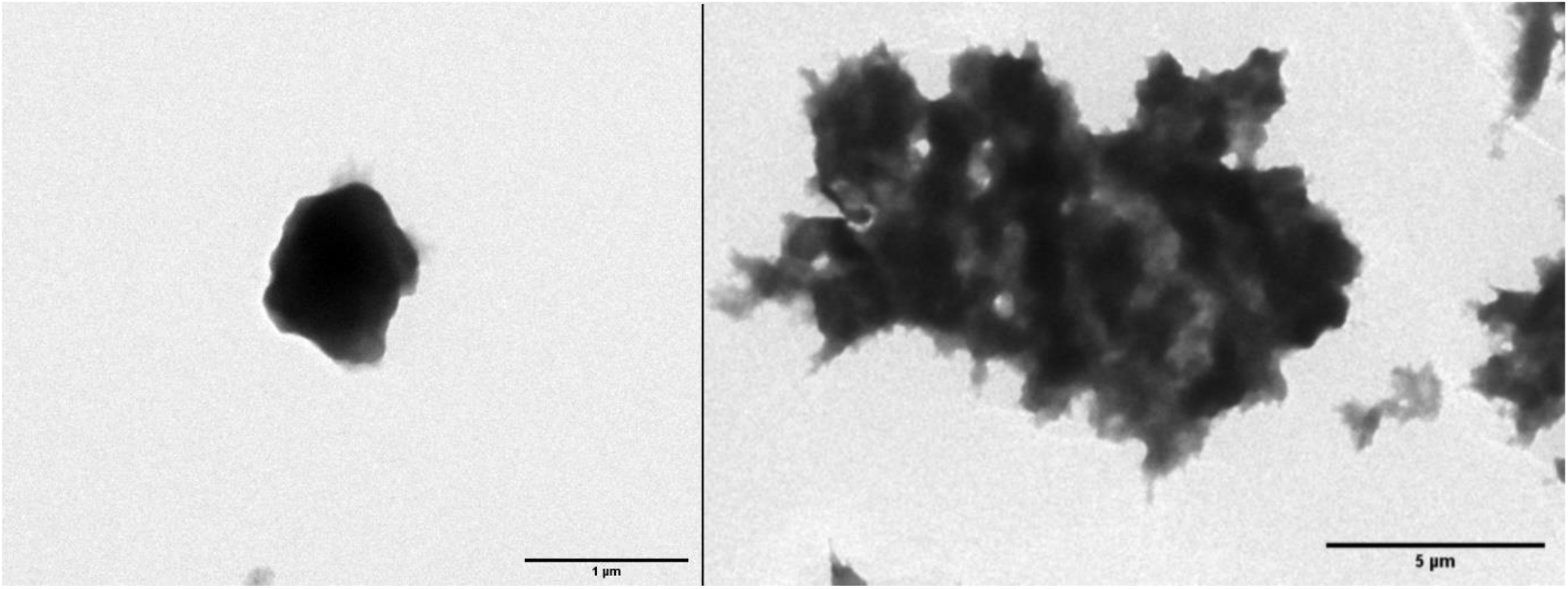
TEM images of a solitary ACC (left) and bulky adducts prepared from ZnCO3 templates. The longest diameter of the solitary ACC measures about 1 μm, while the adduct measures 13.2 μm.

ζ-potential was determined to be −39.4 mV ± 2.94 mV. Each sample was measured three times and the overall mean and standard deviation were calculated.

### Respirometry

Respirometry assays on general functionality were performed in 50 mM sodium phosphate buffer at pH 7.4 and 37 °C (n = 4 each). Interestingly, encapsulated catalase still maintained enzymatic activity. There was a clear dependence of enzymatic activity of ACCs from the volume concentration used for the assay, which is shown in figure 4. This finding is allegeable by the high true activity of the ACCs, leading to experimental set-ups with H_2_O_2_ concentrations out of the ACCs substrate saturation. At higher peroxide concentrations any further observations were impossible due to the formation of oxygen bubbles.

**Figure 4:**
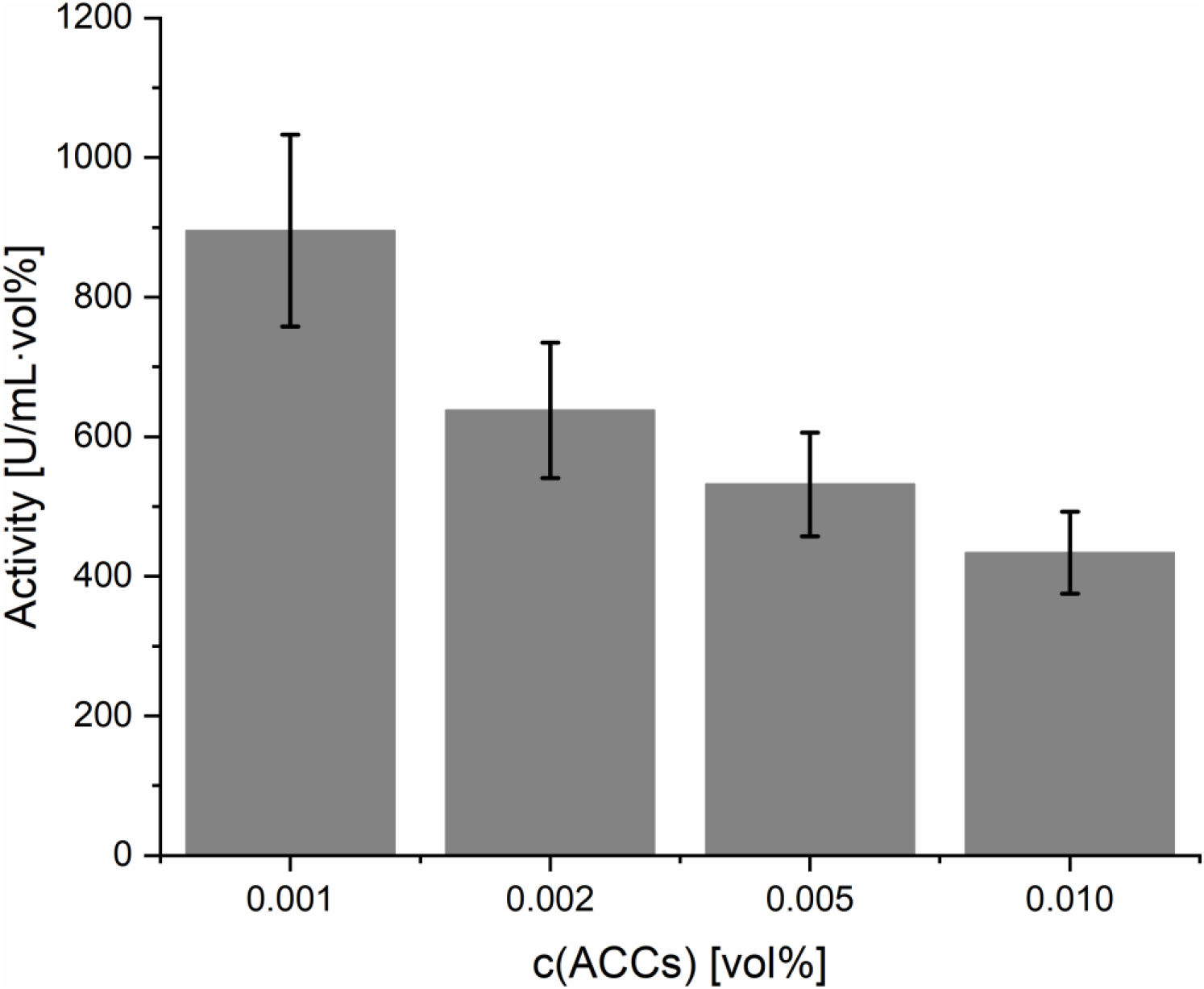
Determination of enzymatic activity of ACCs by respirometry. Mean values are represented by the grey bars. Black bars represent the standard deviation.

Nonetheless, ACCs from MnCO_3_ templates displayed enzymatic activities higher than 1000 U ∙ mL^−1^ ∙ vol%^−1^ in concentrations of 0.001 vol% of the injected suspension.

Due to the smaller volume of ACCs from ZnCO_3_ templates, the amount of catalase used in preparation needed to be more than tripled to reach similar enzymatic activities per volume. At the maximum enzyme-binding capacity of the ZnCO_3_ templates, which equals 7.32-times the amount of enzyme used for MnCO_3_ templates, activities of about 2000 U ∙ mL^−1^ ∙ vol%^−1^ were observed (data not shown).

For better adaption of possible clinical scenarios and application, respirometry assays on storage conditions were performed in AS instead of phosphate buffer (n = 4 each). Furthermore, concentration of injected ACC suspension was decreased to 0.0002 vol%.

ACCs were stored over night at 37 °C, room temperature, 4 °C and −20 °C and compared to the activity of freshly prepared ACCs of the same concentration. For ACCs stored at room temperature and at 37 °C any enzymatic activity of H_2_O_2_ degradation was lost beyond auto-degradation, presumably accelerated by BSA present in AS. ACCs stored at 4 °C still displayed a mean activity of 45.6% with a standard deviation of 4.9% compared to freshly prepared ACCs. ACCs stored at −20 °C even maintained an activity of 67.8% with a standard deviation of 1.5% compared to freshly prepared ACCs. It is therefore tentatively suggested to storage ACCs at −20 °C after preparation. After thawing ACCs should be applied without further delay. If delay is inevitable, thawed ACCs should be stored at 4 °C. The results are depicted in figure 5.

**Figure 5:**
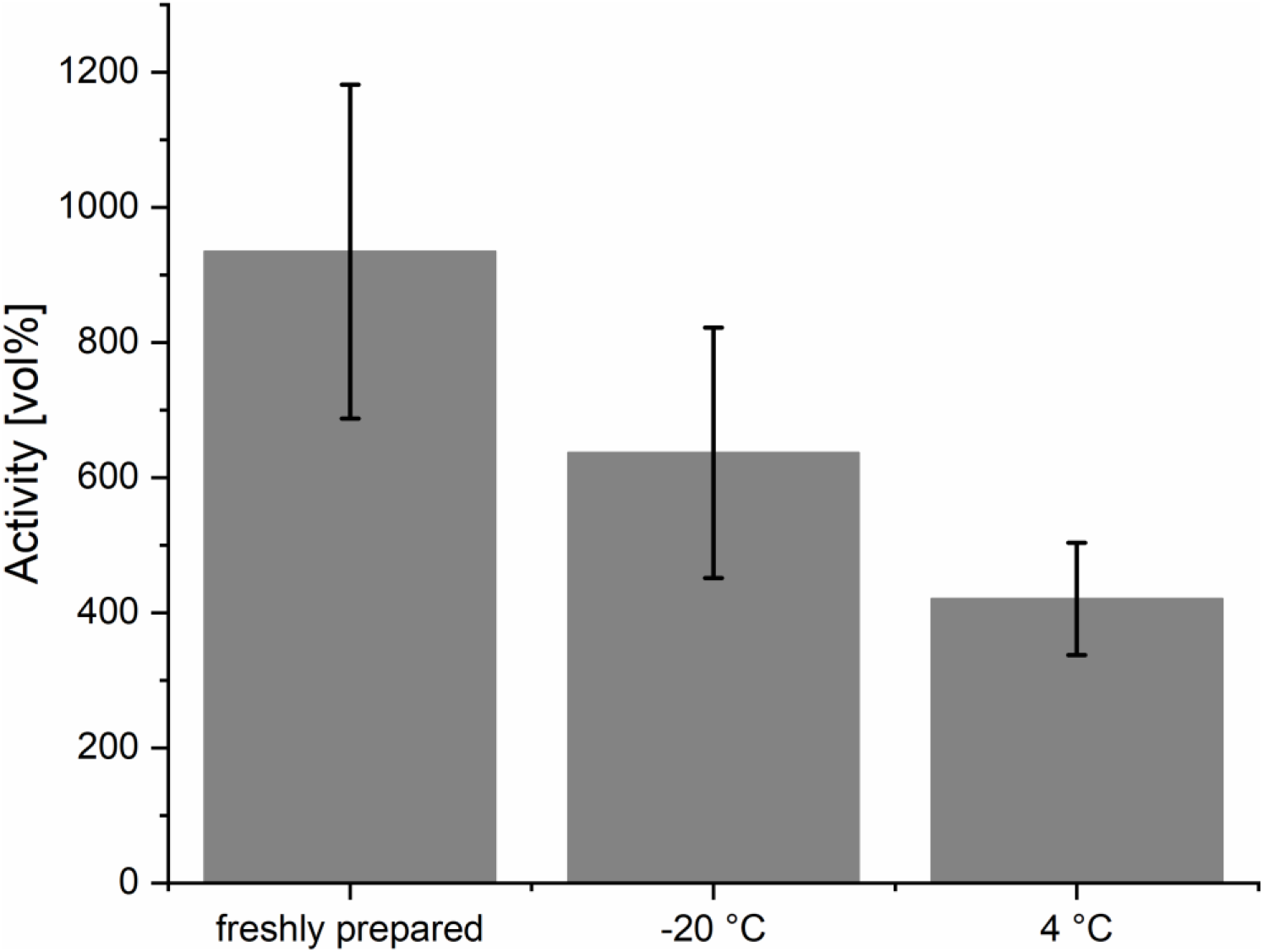
Dependance of ACC activity from storage temperature. Mean values are represented by the grey bars. Black bars represent the standard deviation.

### Cell culture

In the cytotoxicity assay (n = 3), ACCs seemed to be harmless to the L929 cells since uptake of trypan blue did not take place within the observation period of 3 h. Therefore, the observation period was prolonged to 24 h, but again any uptake of trypan blue by L929 cells did not occur.

Protection assays were only performed with ACCs from MnCO_3_ templates under described conditions (*vide supra*) with 50 mM sodium phosphate buffer at pH 7.4 as solvent (n = 3). In line with previous experiments (*vide supra*), ACC concentrations ≥ 0.1 vol% did not evoke any trypan blue uptake in L929 cells, but were evident for ACC concentrations up to 0.05 vol% as shown in figures 6 and 7.

**Figure 6:**
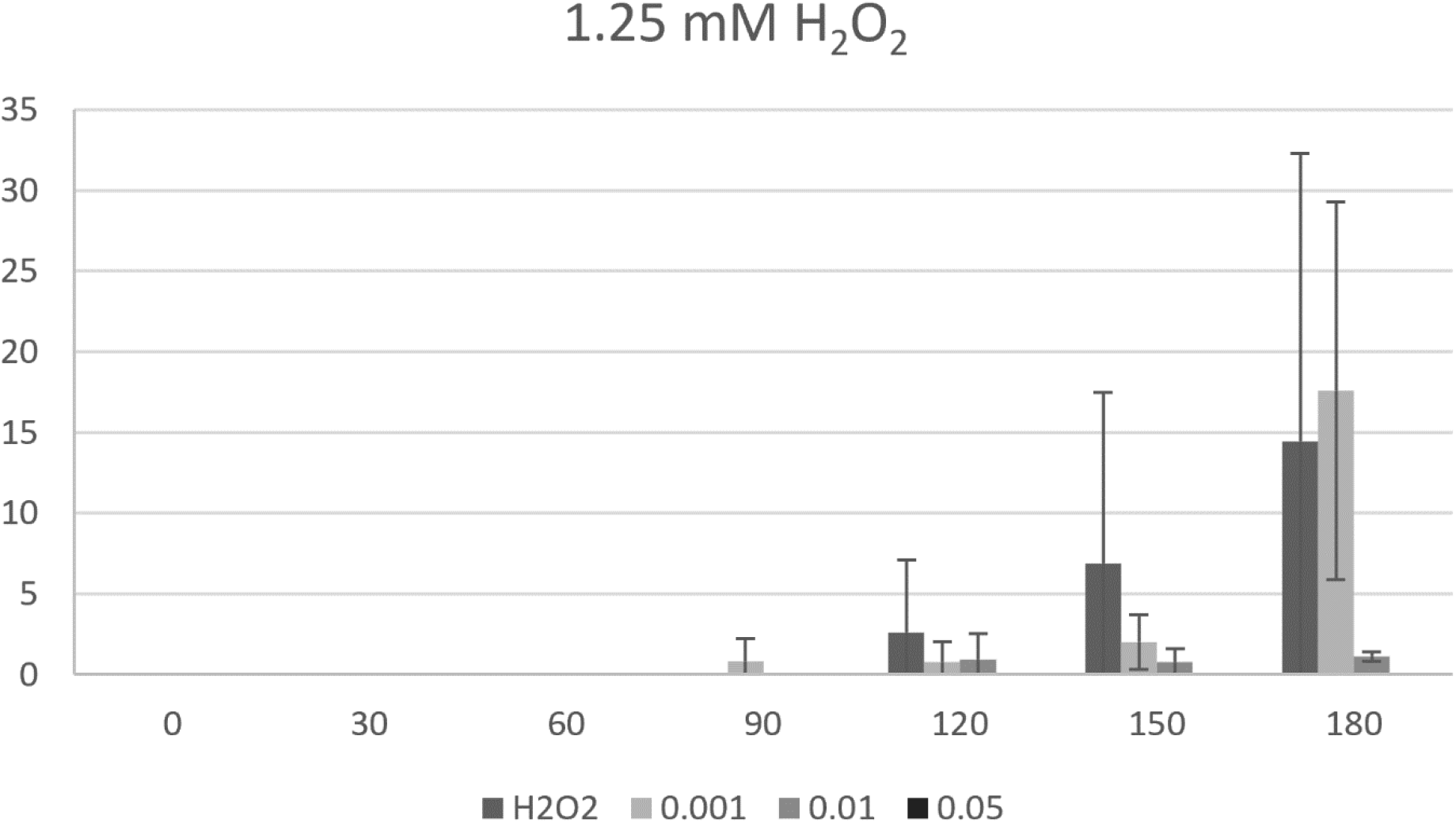
Protection assay performed using low concentrations of H2O2. No protective effect can be observed using low-dose ACCs. The grey bars represent the mean values, while standard deviation is shown by the black bars.

**Figure 7:**
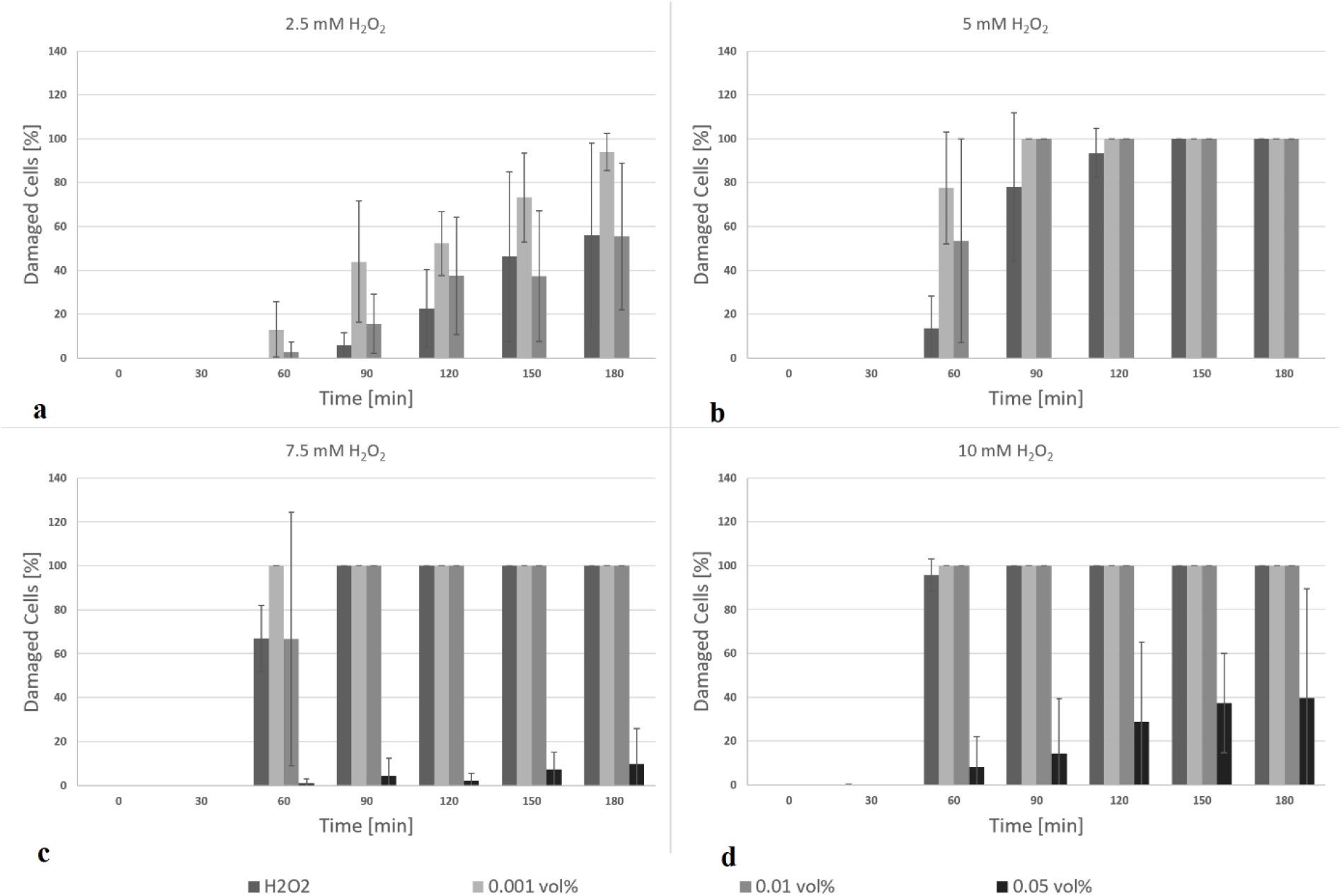
Results of the protection assay using higher peroxide concentrations. The grey bars represent the mean values, while standard deviation is shown by the black bars.

For final H_2_O_2_ concentrations < 2.5 mM a low-dose ACCs (0.001 vol%) application failed in protecting L929 cells (p = 0.34), whereas a medium-dose ACCs (0.01 vol%) already displayed a protective effect (p = 0.08) as seen in figure 6.

Interestingly, for H_2_O_2_ concentrations ≥ 2.5 mM both low-dose and medium-dose ACCs seemed to have an adverse effect, leading to increased cell damage (p < 0.008). High-dose ACCs (0.05 vol%) presented to have significant protective effect for all chosen H_2_O_2_ concentrations. For H_2_O_2_ concentrations < 7.5 mM any trypan blue uptake by L929 cells was prevented by applying ACC concentration of 0.05 vol% or higher. If H_2_O_2_ concentration was ≥ 7.5 mM, only high-dose ACCs displayed significant protective effects (p < 0.005). Figure 7 displays the depicted findings.

## 4 Conclusion

Catalase capsules prepared by the developed method remain highly enzymatically active and decreased peroxide induced cell damage up to 90% at higher concentrations, if dosed correctly. A too low dosage should be avoided because this might result in driving harmful effects. In further assays, co-encapsulation of catalase and other intracellular RBC enzymes, e.g. superoxide dismutase or carbonic anhydrase, could be the next step in developing tools for artificial oxygen carriers. In principle, the presented method can be applied for both perfluorocarbon-based oxygen carriers and hemoglobine-based ones, but, of course, a co-encapsulation of enzymes occurring in erythrocytes together with hemoglobin might be a mandatory formulation in the near future.

## Acknowledgements

The authors thankfully express their gratitude to Mr. Bernd Walkenfort from the Electron Microscopy Unit of the Imaging Centre Essen (IMCES) for the performance of TEM imaging.

## Data availability statement

All data used in this work is publicly available at https://www.doi.org/10.7303/syn23519345

